# Effect of nitrogen on fungal growth efficiency

**DOI:** 10.1101/2020.01.19.911727

**Authors:** D.P. Di Lonardo, A. van der Wal, P. Harkes, W. de Boer

## Abstract

The contribution of fungi to carbon (C) and nitrogen (N) cycling is related to their growth efficiency (amount of biomass produced per unit of substrate utilized). The concentration and availability of N influences the activity and growth efficiency of saprotrophic fungi. When N is scarce in soils, fungi have to invest more energy to obtain soil N, which could result in lower growth efficiencies. Yet, the effect of N on growth efficiencies of individual species of fungi in soil has not been studied extensively. In this study we investigated the influence of different concentrations of mineral N on the growth efficiency of two common soil fungi, *Trichoderma harzanium* and *Mucor hiemalis* in a soil-like environment. We hypothesized that a higher N availability will coincide with higher biomass production and growth efficiency. To test this, we measured fungal biomass production as well as the respiration fluxes in sand microcosms amended with cellobiose and mineral N at different C:N ratios. We found that for both fungal species lower C:N ratios resulted in the highest biomass production as well as the highest growth efficiency. This may imply that when N is applied concurrently with a degradable C source, a higher amount of N will be temporarily immobilized into fungal biomass.

## 1. Introduction

Fungi play a major role in terrestrial decomposition processes and they are important actors in soil organic matter (SOM) dynamics (van der Wal et al. 2013). They are thought to have a relevant contribution to the decomposition of the stable organic carbon (C) pool (Fontaine et al. 2007, 2011). On the other hand, filamentous fungi can promote the formation of soil macroaggregates (Willis et al. 2013) that promotes C sequestration by providing physical protection against decomposers and their degradative enzymes (Wilson et al. 2009).

It is generally assumed that soil microbial communities dominated by fungi have more efficient nitrogen (N) cycling than those dominated by bacteria (Wardle et al. 2004, Van Der Heijden et al. 2008). High N fertilizer additions have been indicated to be the cause of decrease in soil fungal biomass (de Vries et al. 2006, 2007), whereas the cessation of N-fertilizer use can cause a shift from bacterial to fungal dominated systems (de Vries et al. 2006, 2007, Postma-Blaauw et al. 2010). Increases in the abundance of fungi have been linked to a higher efficiency of N cycling and lower N losses from soils (de Vries et al. 2006, 2011, Gordon et al. 2008).

Growth efficiency is defined as the amount of biomass produced per unit of substrate utilized (Sinsabaugh et al. 2013, Mooshammer et al. 2014b). Microbial activity is highest when the C:N ratio of the substrate matches the demands of microbes (Hessen et al. 2004). According to the stoichiometric decomposition theory (Craine et al. 2007), decay processes are driven by the stoichiometry of substrates and adjustment in growth efficiencies may be the most important mechanism to cope with differences between substrate and biomass stoichiometry (Mooshammer et al. 2014b). It is expected that growth efficiencies are influenced by the C:N ratios of the organic substrates. When N is scarce in soils, fungi have to invest more energy in obtaining it, e.g. by producing extra-cellular enzymes, likely resulting in a low growth efficiencies. Generally, knowledge on microbial growth efficiency is of special interest for industrial applications (e.g., to obtain higher biomass or biosynthesized products). However, growth efficiencies have received increased attention of ecologists, due to its important implications for environmental processes (Geyer et al. 2016). Most of the ecological studies focused on soil microbial communities (e.g., Dijkstra et al., 2015; Geyer et al., 2018; Koranda et al., 2014; Mooshammer et al., 2014a; Sinsabaugh et al., 2013). However, fungal biomass production in soil microbial communities is not only affected by fungal growth responses but also by other processes such as predation and competition. Hence, to have a basic understanding of the effect of substrate C:N ratio on fungal growth responses in soil studies with single fungal species are needed.

In this study we investigated the influence of different concentrations of mineral N on the growth efficiency of two common soil fungi, *Trichoderma harzanium* and *Mucor hiemalis* (De Boer et al. 2005, Vinale et al. 2008) in a soil-like environment. As a carbon source we have chosen cellobiose as a model compound for an easily degradable plant-derived carbohydrate (Martínez et al. 2005). We hypothesized that higher nitrogen availability will coincide with higher fungal biomass production and growth efficiency.

## 2. Materials and Methods

Petri dishes (8.5 cm diameter) were filled with 60 g autoclaved, acid -washed quartz sand (granulation 0.1-0.5 mm; Honeywell Specialize Chemicals Seelze GmbH, Seelze, Germany). The lids of the Petri dishes contained butyl rubber stoppers (Rubber BV, Den Haag, The Netherlands) to allow sampling the gas from the headspace of the plates. The sand was amended with 10% (w/w) of a nutrient solution containing (g l^-1^ demineralized water): KH_2_PO_4_, 0.10; K_2_SO_4_, 0.20; Yeast extract (Bacto™; Becton, Dickinson and Company), 0.05; D-(+)-Cellobiose (Sigma-Aldrich), 5.0; MES (2-(N-morpholino) ethanesulfonic acid, Sigma) 5.85. The latter compound was added because acid -washed sand has no buffering capacity. To test the effect of different C:N ratios on fungal growth, the above described nutrient solution received also NH_4_NO_3_ in different amounts. Three nutrient solutions were prepared: i) C-cellobiose:N = 8:1, ii) C-cellobiose:N = 15:1 and iii) C-cellobiose:N = 50:1. The control treatment did not receive any NH_4_NO_3_ addition. The pH of the nutrient solutions was adjusted to 6.5 with NaOH. Before the addition of the nutrient solutions, sand was sterilized by two cycles of autoclaving (30 min at 121 °C, the second one after 24 hours). Next, it was dried at 120 °C for two hours.

Fungal spores (10^4^ spores g sand^-1^) of *Trichoderma harzanium* and *Mucor hiemalis* were obtained from pure cultures (Appendix S1) and mixed with the nutrient-containing sand. In total eight experimental treatments were prepared: *T. harzanium* in sand with no nitrogen (TH No-N), with C:N=8 (TH 8:1), with C:N=15 (TH 15:1) and with C:N=50 (TH 50:1); *M. hiemalis* in sand with no nitrogen (MH No-N), with C:N=8 (MH 8:1), with C:N=15 (MH 15:1) and with C:N=50 (MH 50:1). Each treatment consisted of five replicates. They were sealed with one layer of Diversified Biotech Petri Seal^™^ tape and one layer of Parafilm, to avoid gas exchange with the external environment and maintaining air -tightness. Plates were incubated in the dark at 20 °C.

During the 14-day incubation period, headspace CO_2_ was sampled and measured (Appendix S1). CO_2_ was analyzed at 2, 4, 7, 9, 11 and 14 days. At the end of the incubation period, soil was homogenized by mixing, it was sampled and kept in aliquots in the freezer at −20 °C for ergosterol measurements, DNA extractions and qPCR assays (Appendix S1). We calculated the growth efficiency for each fungus on basis of the amount of ergosterol or ITS copy numbers per amount of CO_2_ released. Growth efficiencies were expressed as relative growth efficiencies where efficiencies of the C:N = 8 treatments were set at 100%.

Differences in respiration, ergosterol, DNA copy numbers and growth efficiencies between treatments were tested with one-way ANOVA followed by post-hoc Tukey’s test, using IBM SPSS Statistics 22. In some cases, due to unequal variances, Tukey’s test was not possible and statistical comparisons were performed by Tamhane’s T2 test. We used linear regression analysis to test the relationship between the different amounts of added N and ergosterol concentrations, and the DNA copy numbers.

## 3. Results and Disccussion

Growth efficiencies for both *M. hiemalis* and *T. harzanium*, as based on ergosterol and DNA copy numbers, showed the same trend, namely an increase with decreasing C:N ratios (Fig. 1A and 1B, P < 0.05). We observed the same pattern for ergosterol production (Fig. S1A, P < 0.05). Furthermore, DNA copy numbers for both fungal species showed an overall increase with an increasing amount of N (Fig. S1B). However, only *M. hiemalis* grown in sand with C:N ratios 8 and 15 had significant higher DNA copy numbers (P < 0.05). On the contrary, respiration fluxes did not increase with decreasing C:N ratios (Fig. S2). Taken together these results indicate that more C-cellobiose was metabolized at lower C:N ratio, implying that not all C-cellobiose has been metabolized in the treatments C:N = 15 and C:N = 50, and certainly not in the control treatments, where there was no addition of mineral N. Our results are in line with our hypothesis, namely that the highest growth efficiency is expected with higher nitrogen availability. A similar growth efficiency pattern was observed for a litter-decomposing fungus grown on maize litter, where the efficiencies increased accordingly with increasing N availability in the plant material (Lashermes et al. 2016). In addition, our results suggest that when N becomes a limiting factor, fungi invest extra energy to obtain N, for instance by recycling their cellular N via controlled autolysis (Santamaria and Reyes 1988) or allocating N to essential metabolic processes (Wicklow 2006).

**Fig. 1:**
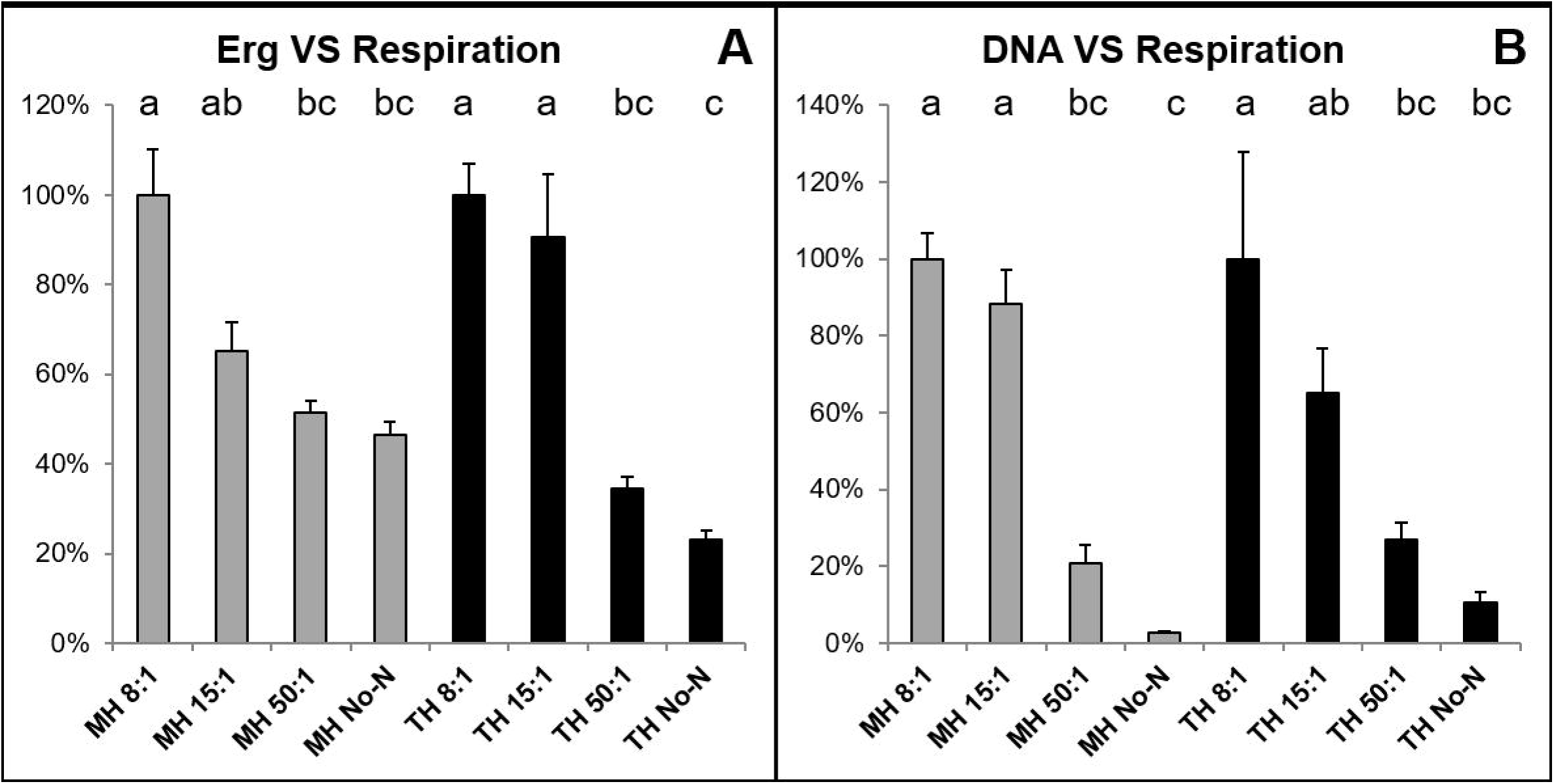
Relative (%) fungal growth efficiencies in sand microcosms containing different C:N ratios after 14 days of incubation. Growth efficiency for each fungus is based on the amount of ergosterol (A) or ITS copy numbers (B) per amount of CO_2_ released. Growth efficiencies of the C:N = 8 treatments were set at 100%. Statistically significant differences (P < 0.05) are indicated with different letters. MH: *M. hiemalis*; TH: *T. harzanium*. No-N: no addition of N; 50:1 is C:N = 50; 15:1 is C:N = 15; 8:1 is C:N = 8. Vertical bars represent standard errors (n = 5).

The linear regression analysis showed a significant positive linear relationship between the amount of added N and ergosterol concentrations (R^2^ = 0.8860; P < 0.0001 and R^2^ = 0.8944; P < 0.0001, for *M. hiemalis* and *T. harzanium* respectively; Fig. S3A) and between the amount of added N and DNA copy numbers (R^2^ = 0.7609; P < 0.0001 and R^2^ = 0.3523; P = 0.006, for *M. hiemalis* and *T. harzanium* respectively; Fig. S3B). Increase of fungal biomass can reduce N losses from soils (de Vries et al. 2006) and, consequently, a higher fungal biomass in soils can be considered as an indicator of higher soil N retention (de Vries et al. 2011). Applications of fertilizers to agricultural soils can result in N losses when crops do not rapidly taken up the added N, and part of the N can be lost via leaching and denitrification (de Vries and Bardgett 2012). Our study indicates that when N is applied concurrently with a degradable C source, a higher amount of N is built into fungal biomass, thereby possibly reducing N losses (Simpson et al. 2007, Liang and Balser 2011). Moreover, our study provides information on how nitrogen influences fungal biomass production and growth efficiency. Knowledge on this effect of N is critical for the assessment of soil C and N budgets.

## Acknowledgments

We are thankful to Jan Geert Bruggink for help in the lab.

## Notes

#### Summary of Updates

Typos were amended

