## Appendix and Supplementary info for "Effect of nitrogen on fungal growth efficiency" (from another version of this preprint: version 1, 2020-01-21)

**Appendix S1**

*Collection of fungal spores*

Fungal spores of *Trichoderma harzanium* and *Mucor hiemalis* were obtained from pure cultures grown on Oatmeal agar (6 g/l Difco, France) and agar powder (2 g/l BOOM CMN, the Netherlands), containing four different antibacterial compounds, namely Chloramphenicol (50 ppm, 12.5 mg/ml final concentration), Streptomycin (50 ppm, 5 mg/ml final concentration), Vancomycin (10 ppm, 2 mg/ml final concentration) and Oxytetracycline (50 ppm, 5 mg/ml final concentration). After 3 weeks of incubation (20ºC in the dark), the spores were washed from the plates with sterile deionized water, collected through Miracloth (Calbiochem, USA) and stored at -80 ºC.

*$CO_2$ measurements*

During the 14-day incubation period, headspace $CO_2$ was sampled through the lid septa and directly injected into 5.9 mL evacuated Exetainer vials (Labco Ltd., Buckinghamshire, UK) and measured using Ultra GC gas chromatograph (Interscience, The Netherlands) equipped with a flame ionization detector (FID) and at Rt-QBOND (30 m, 0.32 mm, ID) capillary column. We used helium as a carrier gas and set the oven temperature at 50 °C. $CO_2$ was analysed at 2, 4, 7, 9, 11 and 14 days. After each measurement the Petri dishes were opened under the flow cabinet for aeration and sealed again after 15 minutes.

*Ergosterol measurements*

Ergosterol, a sterol found in fungal membranes, was used as a biomarker for fungal biomass in the sand (de Ridder-Duine et al. 2006). Briefly, 4 g of moist sand was shaken with 6 ml of methanol in the presence of glass beads, to disrupt the fungal mycelium and to release the ergosterol into the extractant. After centrifugation and filtration, ergosterol was measured on a 1260 Bio-inert LC coupled with a 6460 QQQ (Agilent, Santa Clara, USA).

*DNA extraction and qPCR assay*

DNA was extracted from sand using the PowerSoil®DNA Isolation Kit (MOBIO Laboratories, Carlsbad, California, USA) according to the manufacturer's instructions with some modifications: after adding solution C1 (causing cell lysis), samples were incubated at 60 °C for 30 min; solutions C2 and C3 (for the removal of contaminating inorganic and organic matter) were added (100 µl each) to the samples; after adding solution C6 (releasing DNA from spin filter), samples were incubated at 30 °C for 10 min. DNA was then collected in Eppendorf tubes and stored at -20 °C.

To determine the amount of DNA present in the samples, quantitative PCR assays were performed using different primer combinations based on taxon-characteristic internal transcribed spacer (ITS) regions (Table S1). The total end volume of the qPCR assays was 20 µl containing: 3 µl of 100 times diluted DNA, 1 µl of each primer (final concentration: 10ng/ µl), 5 µl of Milli-Q water and 10 µl iQ SYBR Green Supermix (BioRad). The following temperature profile was run on a Biorad iQ5 thermal cycler: 95°C for 3 min followed by 40 x (95°C, 30 s; 60 °C, 30 s; 72 °C, 30 s). To establish the relationship between Ct-values and the concentration of target fungal DNA (ng/ µl), a calibration curve was made for each primer combination using a dilution series of the corresponding DNA as a template. The concentration of DNA was measured using a NanoDrop ND-1000 UV–VIS spectrophotometer (NanoDrop Technologies).

**Table S1:** Primers used for the qPCR assay. Fungal primers are based on taxon-characteristic internal transcribed spacer (ITS) regions.

| Target | | Primer sequence (5'-3')<br>F: forward, R: reverse | Reference | $T_a$ | Parameters defining the linear relationship between $C_t$ and DNA concentration ($C_t$ = a x $^{10}$log [concentration] + b) | | |
|---|---|---|---|---|---|---|---|
| | | | | | a | b | $R^2$ |
| **Ascomycota** | F | GGAAGTAAAAGTCGTAACAAGG | Nikolcheva et al., 2005; White et al., 1990 | 60 °C | -3.04 | 18.73 | 0.981 |
| | R | CGTTACTRRGGCAATCCCTGTTG | | | | | |
| **Zygomycota (Mucorales)** | F | CAGTTAAAACGTCCGTAGTCAA | Harkes et al., 2017 | 60 °C | -4.55 | 14.56 | 0.988 |
| | R | CACCACCACCCATAGAATCTA | | | | | |

**Supplementary information**

**Fig. S1:** Concentrations of ergosterol (A) and ITS rDNA copy numbers (B) of *M. hiemalis* and *T. harzianum* grown for 14 days in sand microcosms at different C:N ratios. Statistically significant differences ($P < 0.05$) are indicated with different letters. MH: *M. hiemalis*; TH: *T. harzanium*. No-N: no addition of N; 50:1 is C:N = 50; 15:1 is C:N = 15; 8:1 is C:N = 8. Vertical bars represent standard errors ($n = 5$).

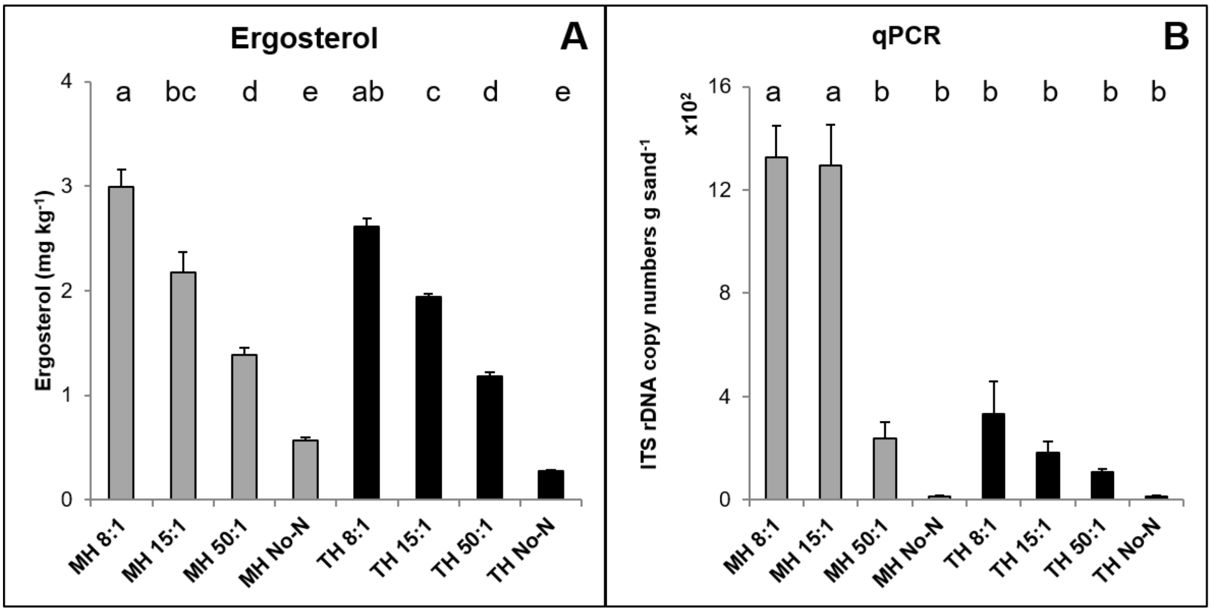

**Fig. S2:** A) Accumulation dynamics of $CO_2$ during 14 days of incubation of *M. hiemalis* and *T. harzianum* in sand microcosms at different C:N ratios; B) Total accumulated $CO_2$ concentrations after 14 days of incubation. Statistically significant differences ($P < 0.05$) between treatments are indicated with different letters. MH: *M. hiemalis*; TH: *T. harzanium*. No-N: no addition of N; 50:1 is C:N = 50; 15:1 is C:N = 15; 8:1 is C:N = 8. Vertical bars represent standard errors (n = 5).

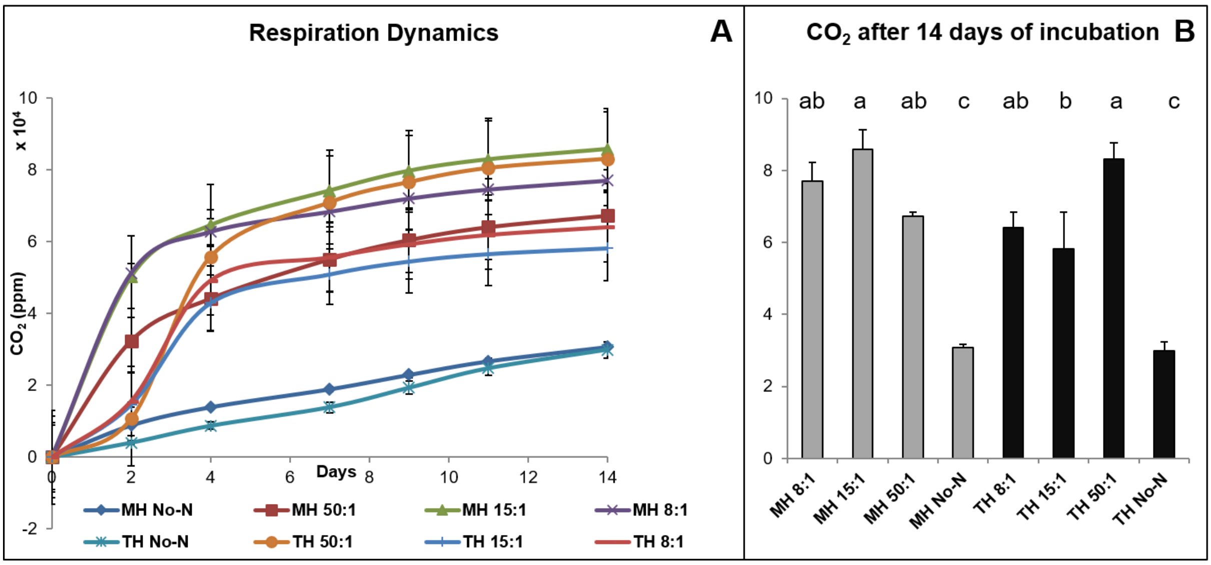

**Fig. S3:** Linear regression analysis between the different amounts of added ammonium nitrate ($NH_4NO_3$) and concentrations of ergosterol (A), and ITS rDNA copy numbers (B) of *M. hiemalis* and *T. harzianum* grown for 14 days in sand microcosms.

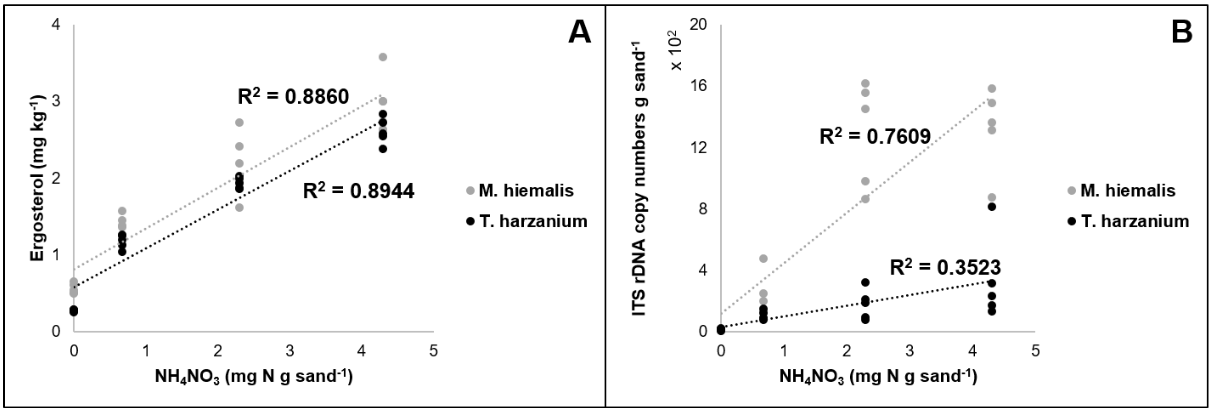
